# High-risk clonal groups of Avian Pathogenic *Escherichia coli* (APEC) demonstrate heterogeneous phenotypic characteristics *in vitro* and *in vivo*

**DOI:** 10.1101/2025.04.10.648166

**Authors:** James R. G. Adams, Huijun Long, Charlotte A. Birdsall, Kamran Qureshi, Emma King, Sara Perez, Keith Warner, Shahriar Behboudi, Roberto M. La Ragione, Jai W. Mehat

**Affiliations:** Department of Comparative Biomedical Sciences, School of Veterinary Medicine, Faculty of Health and Medical Sciences, University of Surrey, Guildford, Surrey, GU2 7AL, United Kingdom; The Pirbright Institute, Pirbright, Woking, Surrey, GU24 0NE,United Kingdom; Discipline of Microbes, Infection and Immunity, School of Biosciences, Faculty of Health and Medical Sciences, University of Surrey, Guildford, Surrey, GU2 7XH, United Kingdom; Poulty Health services, Sheriff Hutton, York, YO60 6RZ, United Kingdom; Avara Foods Ltd, 1 Willow Road, Brackley, Northamptonshire, United Kingdom, NN13 7EX; Bristol Veterinary School, University of Bristol, Langford, BS40 5DU, United Kingdom

**Author notes:** **Corresponding author** Jai W. Mehat.

**Keywords:** APEC, avian pathogenic *Escherichia coli*, sequence types, cell culture, *Galleria mellonella*

## Abstract

Avian Pathogenic *Escherichia coli* (APEC), a major bacterial pathogen of poultry, is comprised of a diverse range of high-risk clonal groups. However, phenotypic interactions with the avian host cell and how they may differ between lineages remains poorly understood. Therefore, the ability of predominant and outbreak-associated APEC clonal groups to invade and survive within avian host cells, as well as virulence within the *Galleria mellonella* infection model was investigated. The molecular characterisation of APEC isolated from an outbreak of colibacillosis in turkey poults in the UK, identified APEC sequence type (ST)-101 as the dominant clonal group, carrying a high number of virulence factors. As such, ST-101 was compared as an outbreak-associated lineage to a range of predominant APEC high-risk clonal groups (ST-23, ST-140, ST-95, ST-117). Utilising *in vitro* cell culture models, APEC isolates displayed comparable adhesion to 8E11 chicken epithelial gut and HD11 chicken macrophage cell lines. However, a trend of increased invasion of the 8E11 cells, and intracellular survival within HD11 macrophages was observed for ST-95, ST-101, and ST-140 APEC, relative to ST-23 and ST-117, suggestive of pronounced phenotypic differences between clonal groups. However, in HD11 cell assays, no difference in magnitude of elicited immune response was observed between lineages, indicating lineages had differing capacities to resist phagocyte killing. *In vivo* virulence in the *Galleria mellonella* infection model was also observed to differ between APEC genotypes, with ST-117 inducing the highest mortality, despite the comparatively lower epithelial invasion and intramacrophage survival to other lineages. Collectively, this suggests a distinct phenotypic profile associated with high-risk clonal groups within APEC, potentially allowing the future development of broad-spectrum disease management strategies.

## Introduction

The vast majority of *E. coli* are commensal organisms of the vertebrate gut, exhibiting a mutualistic relationship with its host, preventing pathogen colonisation by competitive exclusion (1), priming the immune system against infection (2, 3), and producing vitamins for the host (4). However, a minority of genetic lineages are responsible for a variety of extra-intestinal pathologies. Pathogenic, or pathobiont lineages of *E. coli*, may arise through the horizontal acquisition of virulence determinants, which are often independent events in disparate phylogenetic backgrounds. This has resulted in the repeated emergence of distinct pathogenic lineages capable of eliciting disease in healthy hosts (5, 6). *E. coli* lineages are frequently categorised by multi-locus sequence typing (MLST) and as distinct sequence types (STs) and clonal groups. Primary pathogenic lineages are the most common in humans and are broadly divided into intestinal pathogenic *E. coli* (InPEC) (STc11, STc29, ST16 and ST17) and extra-intestinal pathogenic *E. coli* (ExPEC) (STc131, STc95, STc73, and STc69) pathotypes (6). However, avian species regularly experience primary and opportunistic secondary infection, with the causative *E. coli* defined as belonging to the avian pathogenic *E. coli* (APEC) pathotype (7-10). APEC is the causative agent of avian colibacillosis, a global financial, and animal health and welfare issue which results in increased bird mortality, diminished egg production, and reduced hatchability, affecting every production system in all farmed species of poultry (11).

Classification of *E. coli* as APEC has traditionally been based on isolation from a diseased avian host, leading to a wide number of phenotypically and genotypically disparate organisms being termed collectively (12). PCR-based screening of presumed APEC isolates has been suggested, with several frameworks proposed for the determination of APEC based on the presence of a combination of virulence factors. However, this approach is limited in its capability to correctly distinguish commensal or pathogenic *E. coli* isolated chickens (12-14). Multiple sequence types are associated with APEC globally with the STc-23, ST-95, ST-117, ST-140, and ST-428/429 high-risk clonal groups being most associated with avian colibacillosis (12, 15-17). Multiple strategies for the control of APEC have been developed, including vaccination aimed at targeting the globally predominant serotype O78 comprised of the STc-23 and ST-117 lineages (12) However, heterologous protection from vaccination can be inconsistent (18-22), potentially allowing novel APEC lineages to circumvent vaccine protection.

Plasmids can facilitate the rapid dissemination of genes within microbial environments, impacting bacterial fitness (23). The Colicin V (ColV) plasmid is over-represented in APEC (24, 25), and encodes a number of virulence related genes such as adhesins, toxins or iron acquisition systems (26, 27). ColV/IncF plasmids are widespread in *E. coli*, having been detected in 98% and 73% of septicaemic chickens and turkeys isolates, respectively (28). Considering epidemiological studies implicate specific STs in the majority of outbreaks, it is clear that these virulence plasmids work in concert with defined phylogenetic backgrounds to mediate disease. However, the exact contribution of ColV to pathogenicity and the relative importance of these plasmids in the context of differing *E. coli* genetic backgrounds is not fully understood (13, 29). This highlights the need to unravel the precise genetic determinants which contribute to APEC virulence and elucidate how these may differ between high-risk APEC clonal groups.

This study aimed to characterise the interaction of APEC with avian epithelial cells and macrophages. In this study, we identify the *E. coli* ST-101 sequence type as a highly prevalent lineage implicated in a recent colibacillosis outbreak in turkey poults. Comparison of the pathogenic potential of ST-101 relative to APEC isolates belonging to the predominant, high-risk clonal groups demonstrated that APEC clonal types differ in their capacity to survive intracellularly in avian macrophages and epithelial cells. Finally, using the *in vivo Galleria mellonella* infection model we demonstrate that virulence capacity is variable between APEC sequence types. Our results demonstrate remarkable phenotypic variation between APEC groups and highlight the need for intervention strategies that can simultaneously target a broad range of pathogenic lineages.

## Methods

### Bacterial isolates and DNA extraction

Bacterial isolate collection received a favourable ethical opinion by the University of Surrey’s animal ethics committee. A total of 91 *E. coli* isolates were collected from the heart, liver, or caeca of found dead or euthanised turkeys housed on a commercial UK farm which were clinically confirmed to have colibacillosis. Isolates were streaked onto MacConkey Agar No. 3 (Oxoid, Basingstoke, UK) to confirm purity, prior to selection of a representative colony and preparation of a bacterial stock in Pro-Lab Diagnostics Microbank tubes (Fisher, Basingstoke UK) for storage at -80°C. For the use in assays, *E. coli* isolates were streaked onto MacConkey Agar No. 3 (Oxoid, Basingstoke, UK) while *S*. Typhimurium SL1344 was streaked onto nutrient agar (Oxoid, Basingstoke, UK), before incubation at 37°C, aerobically for 18 hours. For broth cultures, a single representative colony of each isolate was used to inoculate 10 mL of fresh LB broth in a sterile 50 mL Falcon tube before aerobic incubation at 37°C with shaking at 225 RPM, for 18 hours. From prepared cultures, DNA was extracted from a broth culture of each isolate, using a Wizard Genomic DNA purification kit (Promega, Wisconsin, United States) following the manufacturer’s instructions, before quantification using a Biodrop spectrophotometer (DKSH, Zurich, Switzerland) and storage at -20°C. Extracted DNA was sent to MicrobesNG (Birmingham, UK) to undergo Illumina MiSeq sequencing (Table S1).

In addition, previously characterised APEC isolates and *Salmonella* Typhimurium SL1344 held in the Surrey Animal Pathogen (SAP) collection, in Pro-Lab Diagnostics Microbank tubes, were also used within this study (Table 1). Selection of APEC lineages for inclusion within the panel was based on previously published literature on APEC prevalence (12, 15, 30-34).

**Table 1:**
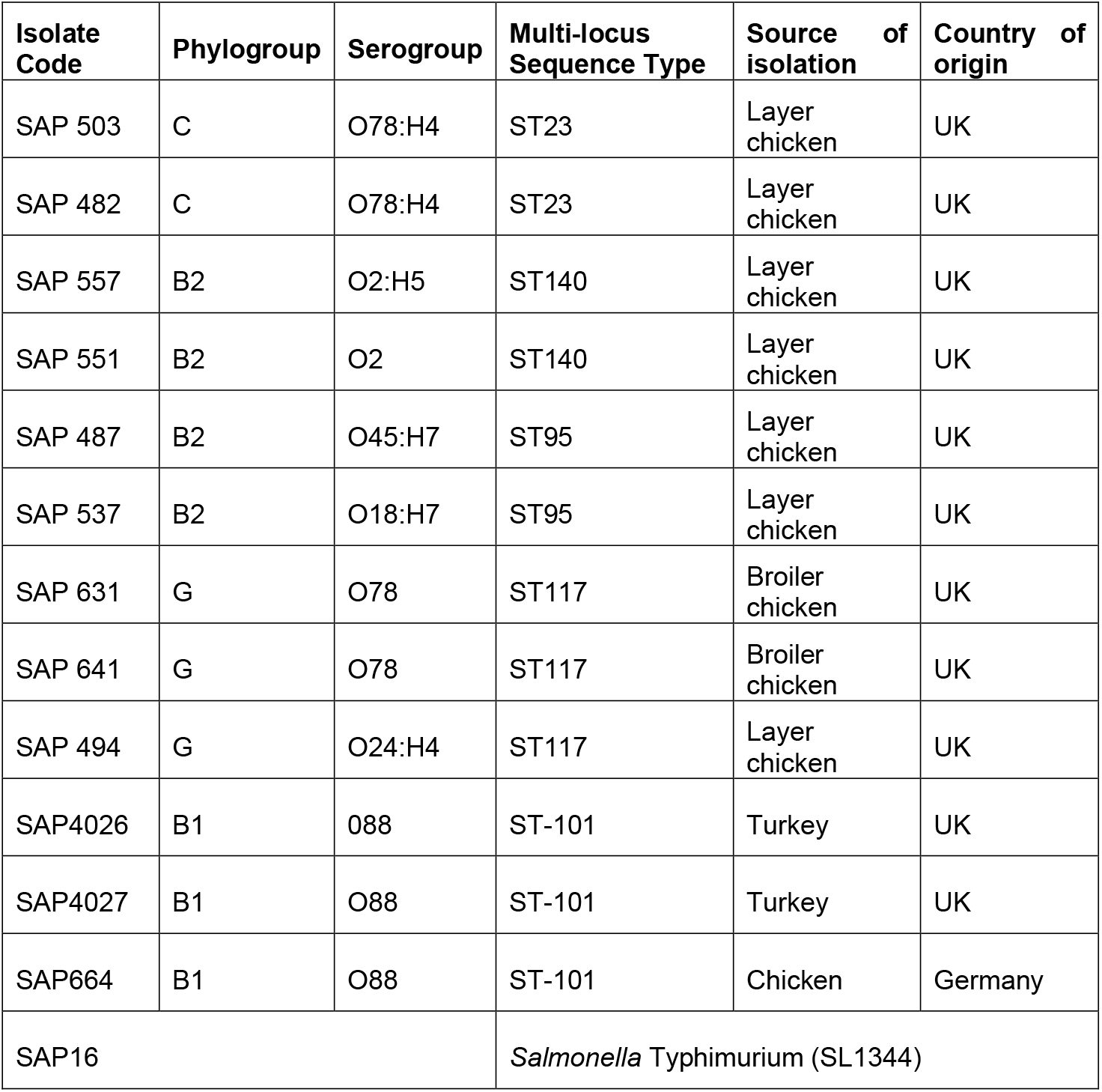
List of bacterial isolates identified used in this study for cell culture assays. All isolates were whole genome sequenced, and this data used to classify isolates according to Phylogenetic group, serotype, and multi-locus sequence type.

### Genomic assembly, annotation, phylogenetic reconstruction, and typing

Following MiSeq sequencing, raw reads were assembled into contigs using Shovill (https://github.com/tseemann/shovill), followed by annotation using Prokka (https://github.com/tseemann/prokka). The 91 total genome sequences were used to construct a phylogenetic tree through identification of the core genomes and core single nucleotide polymorphisms (SNPs) using Parsnp v1.2 software (https://github.com/marbl/parsnp) and visualised using iTOL (https://itol.embl.de/). Phylogroup and serotype was determined using the EcOH database (35). The multi-locus sequence type (MLST) of APEC genomes was determined using the MLST software (Seemann T, mlst, Github https://github.com/tseemann/mlst) with allelic variation within the sequences of seven *E. coli* housekeeping genes (Warwick scheme); *adk, fumC, gyrB, icd, mdh, purA*, and *recA*, used to determine sequence type (36).

### Growth and maintenance of cell lines

The chicken gut epithelial cell line 8E11 and chicken macrophage-like cell line HD11 were used within this study. 8E11 cells were maintained in Dulbecco’s Modified Eagle Medium/Nutrient Mixture (DMEM) F-12 + GlaMAX (ThermoFisher, Paisley, UK) with 10% foetal bovine serum (FBS) (ThermoFisher, Paisley, UK) and 1% Penicillin/ Streptomycin (ThermoFisher, Paisley, UK) at 37°C + 5% CO^2^. The HD11 cell line was maintained in Roswell Park Memorial Institute (RPMI) 1640 Medium (ThermoFisher, Paisley, UK) with 5% FBS, 5% chicken serum (CS) (ThermoFisher, Paisley, UK) and 1% Penicillin/ Streptomycin at 37°C + 5% CO_2_. When 80% confluency was reached, cells were removed from flask using 0.25% trypisin (ThermoFisher, Basingstoke, UK) and used to seed 24 well plates (Greiner, Stonehouse, UK) at a concentration of 2 × 10^5^ cells/ml.

### Determination of bacterial intracellular survival

8E11 or HD11 cells were seeded in duplicate 24 well tissue culture treated plates at 2 × 10^5^ cell/mL in a total of 1 mL media per well at least 24 hours prior to assay to ensure adherence. Concurrently, a single representative colony was used to inoculate 10 mL of fresh LB broth in a 50 mL Falcon tube before aerobic incubation at 37°C with shaking at 225 RPM, for 18 hours. Following incubation, 100 μL of bacterial culture was removed and used to inoculate 9.9 mL LB broth or 9.9 mL LB broth in a 50 mL centrifuge tube before growth for two hours at 37°C, aerobically with shaking at 225 RPM. Triplicate wells in duplicate plates were challenged with a MOI of 100 (8E11) or 10 (HD11) of APEC culture before centrifugation at 4°C for 3 minutes at 300 x g. Plates were incubated for two hours before aspiration of the media and washing of the plates twice with PBS. To determine association media was removed and cells were washed twice with PBS before 1 ml of 1% Triton 100X (Merck, Kenilworth, UK) was added to the wells. The solution was gently pipetted to disrupt the eukaryotic cell membrane and resuspend the associated bacteria. The CFU/ ml of the inoculant was calculated using the Miles and Misra technique (37). To determine intracellular survival, of 1 mL cell culture media supplemented with 100 µg/mL gentamicin and incubation for an additional 2, 4, or 16 hours. Cell supernatant was then aspirated and HD11 were washed twice with warm PBS and 1 mL 1% triton X in PBS was added to lyse cells. Bacterial viability within the cell lysate was then quantified using the Miles and Misra technique (37).

### Determination of nitric oxide and reactive oxygen species production following bacterial challenge

HD11 cells were seeded in 24 well tissue culture treated plates at 2 × 10^5^ cell/mL in a total of 1 mL media per well at least 24 hours prior to assay to ensure adherence. Triplicate wells were challenged with bacterial isolates at a MOI of 10 were centrifuged at 4°C for 3 minutes at 300 x g before incubation at 41°C + 5% CO_2_ for 1 hour.

To determine nitric oxide (NO) production cell supernatant was removed and replaced with 1 mL of RPMI + 5% FBS + 5% CS + 100 µg/ mL gentamicin and incubated for a further four hours. The cell supernatant was transferred to a 1.5 mL microcentrifuge tubes and centrifuged at 2500 x g for 10 minutes at 4°C to pellet cell debris. Following this, 150 µL of the supernatant was mixed with 130 µL ddH_2_O and 20 µL Griess reagent (ThermoFisher, Paisley, UK) in triplicate in a 96 well plate (Greiner, Stonehouse, UK). The plate was incubated at room temperature, protected from light for 30 minutes and absorbance was read at 548 nm on a TECAN Spark plate reader and nitrate concentration (the autooxidation product of NO) was determined using a calibration curve.

To determine ROS generation supernatant was removed and replaced with 1 mL of RPMI + 5% FBS + 5% CS + 100 µg/ mL gentamicin + 20 µM DCFA (D399, ThermoFisher, Paisley, UK). The plate was incubated in the Clariostar plate reader (BMG Labtech, Ortenberg, Germany) at 41°C + 5% CO_2_ for a further ten hours with fluorescence at 495 nm excitation and 529 nm emission measured every 15 minutes.

### *Galleria mellonella* infection model

Larvae in the fifth or sixth instar stage were purchased from Livefoods UK (Sheffield, UK), maintained at 17°C on woodchips and used in assays within two weeks of delivery. Larvae were handled using ethanol sterilised blunt-nose forceps and separated into groups of suitable larvae based on uniform colouring and ability to right themselves after being inverted and divided into groups of 10 in duplicate. APEC isolates (Table 1) were streaked onto MacConkey agar plates and incubated aerobically at 37°C for 18 hours before colonies were suspended in PBS at a OD_600_ of 0.125 before diluting to reach a final concentration of 1×10^3^ CFU per 10 µL. Larvae were then infected as described previously (38, 39), in the top right proleg with either 10 µL of bacterial suspension or 10 µL sterile PBS using a Hamilton 26S Microliter syringe (Merck, Kenilworth, UK). Following injection, larvae were placed on 90-mm filter paper in an inverted Petri-dish and incubated for 72 hours at 37°C. Mortality was determined every 24 hours by observing melanisation and ability to respond to physical stimulation (38).

## Results

### APEC ST-101 are the most prevalent sequence-type implicated in a colibacillosis outbreak and encode a high density of APEC-associated virulence determinants

An outbreak of colibacillosis in turkey poults was characterised by whole genome sequencing (WGS). APEC isolates recovered from turkey poults at *post-mortem* examination were determined to belong to 18 distinct sequence types (STs), with ST-101 observed to be dominant within the outbreak (26.3% of all isolates) (Figure 1). Isolates belonging to ST-7804 were second most prevalent, accounting for 15.3% of all APEC recovered. Comparison of metadata also revealed an association between disease state and sequence type. Of the 24 isolates belonging to ST-101, 23/24 (96%) were recovered from sick birds and 20/24 (83%) from birds which died during rearing. Meanwhile, only 4/14 (28%) isolates belonging to ST-7804 were isolated from diseased or dead birds.

**Figure 1:**
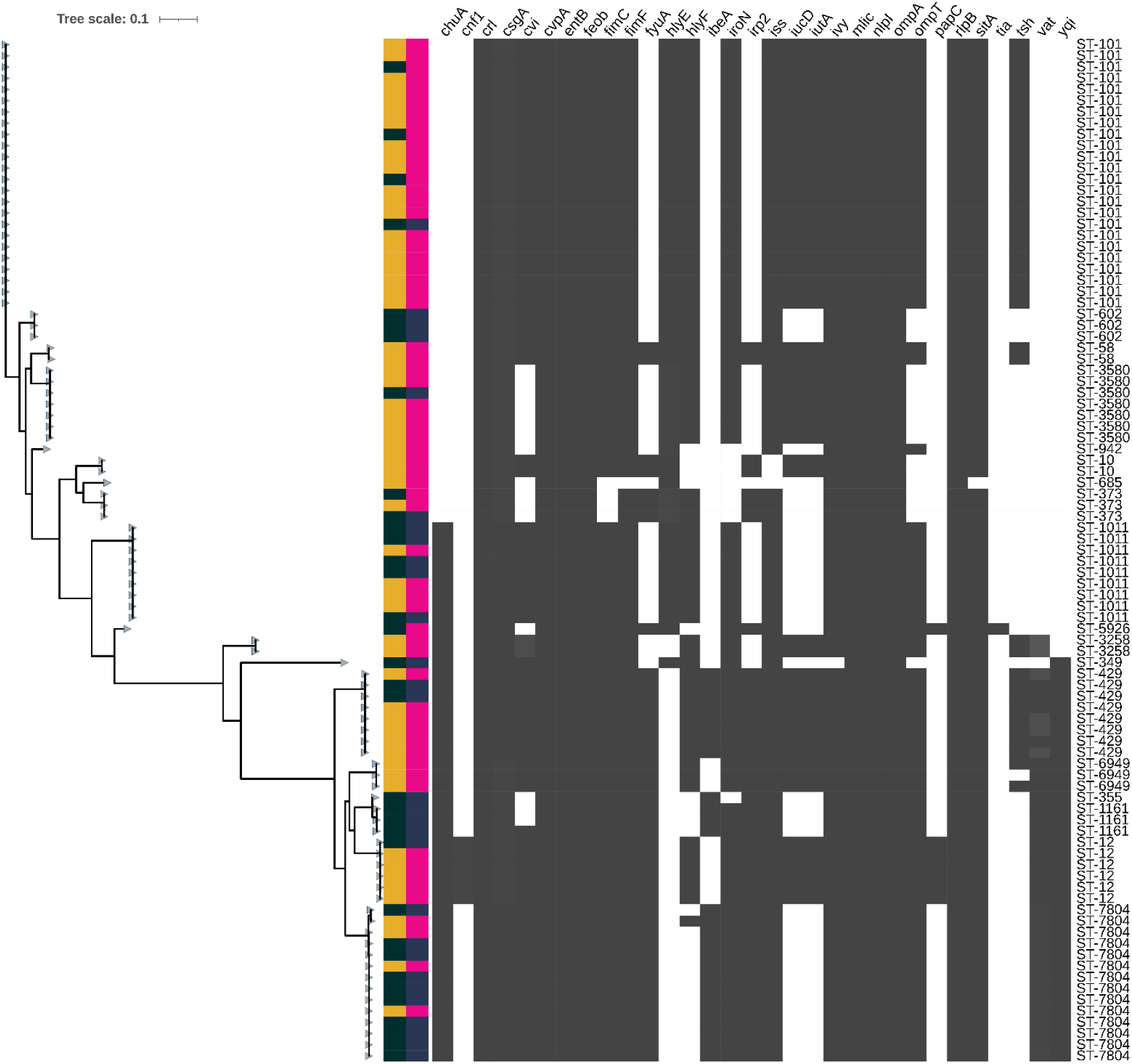
Phylogenetic reconstruction of 91 *E. coli* genomes (Table S1) recovered from a colibacillosis outbreak in turkey poults. Whole genome sequences of *E. coli* isolated from the heart, liver, and caeca of turkeys suspected of colibacillosis were used to reconstruct core-genome phylogeny using ParSNP (42). Health-associated metadata of bird fate (dead or culled) and health status (sick/moribund or healthy) denoted by the colour strip to the right of the cladogram. Presence of encoded APEC-associated virulence genes determined using a custom Abricate (https://github.com/tseemann/abricate) database using an 80% minimum sequence identity threshold. The presence or absence of a virulence gene within the bacterial genome is denoted by a grey box in the heatmap (grey=present, white=absent). Sequence type (ST) was established using Warwick classification scheme (Seemann T, *mlst*, Github https://github.com/tseemann/mlst) and denoted to the right of the heatmap

Due to its dominance within the outbreak and association with bird mortality and morbidity, ST-101 APEC were characterised further. ST-101 APEC were determined to belong to phylogroup B1 and encode the O88:H8 serotype antigens. *In silico* screening of virulence genes revealed a highly conserved virulence profile, encoding genes associated with colonisation and adhesion (*crl, csgA, fimC*, and *fimF*), iron acquisition systems (*iutA, iucC, iucD*, and *sitA*), protectins (*iss, mliC, ompT*, and *OmpA*), and toxins (*hlyE, hlyF*, and *vat*) (40). To investigate the distribution of these genes between the chromosome and plasmid of the ST-101 isolates hybrid short and long read sequencing of an isolate designated as SAP4026 was performed. Five closed contigs were identified, consisting of the bacterial chromosome and four plasmids. One plasmid identified as a Colicin V plasmid encoded a large density of virulence associated genes (Figure 2). This included a range of iron acquisition genes including *iroBCDEN, sitABC, iucABCD*, and *iutA*. Colicin A, B, C and M genes were also identified on the same plasmid, alongside *tsh, iss*, outer membrane protease family protein *ompT*, and antimicrobial peptide resistance gene *Mig-14*. A range of conjugation machinery was also identified within the plasmid, suggesting a capacity for intra- and inter-species horizontal transfer.

**Figure 2.**
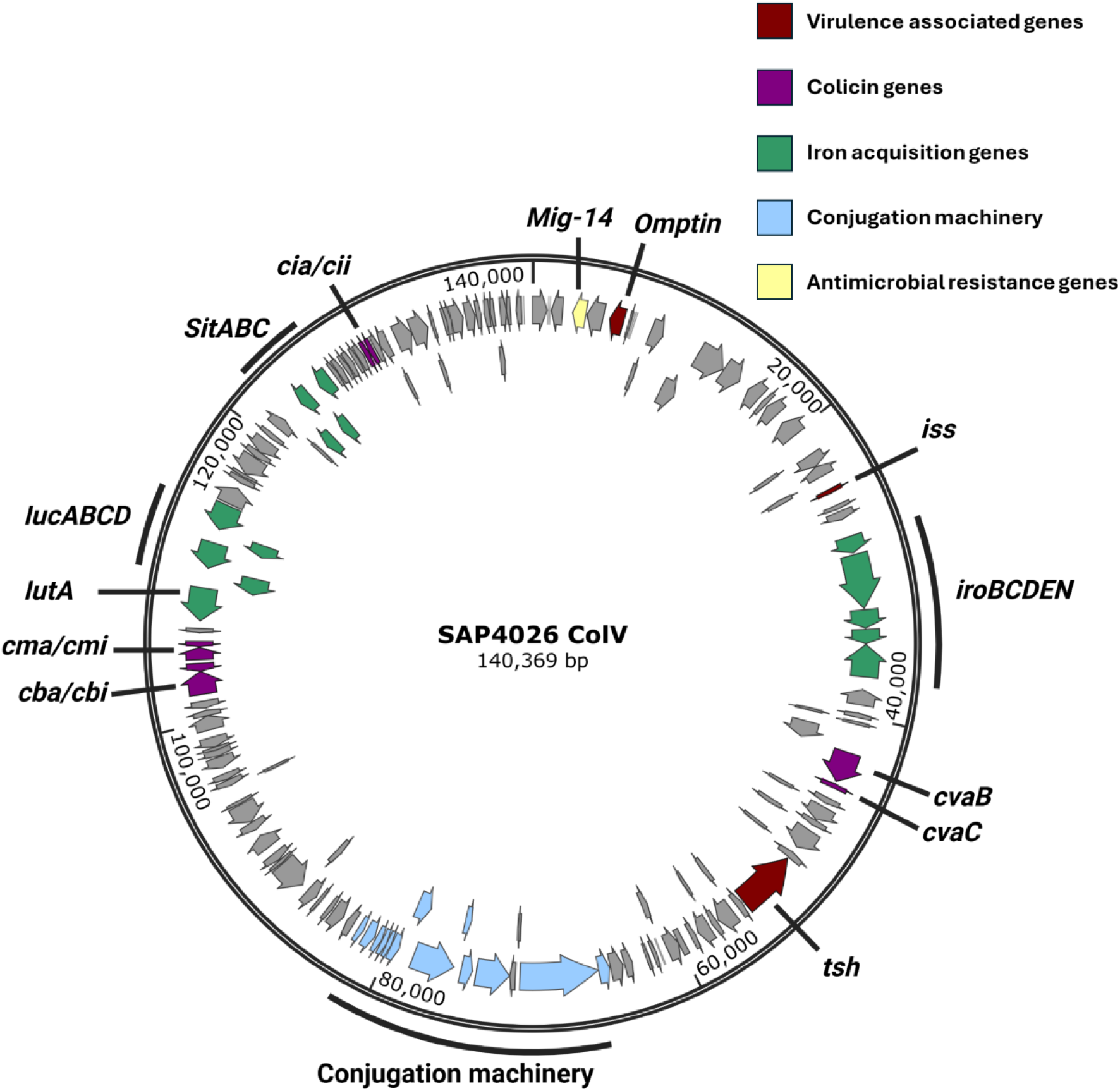
Plasmid map of ColV plasmid harboured by ST-101 SAP 4026. Genes identified by Bakta (https://github.com/oschwengers/bakta) annotation of circular plasmid contig determined through hybrid sequencing. Red = virulence associated genes. Purple = colicin genes. Green = iron acquisition genes. Blue = conjugation machinery. Yellow = antimicrobial resistance genes.

To determine whether the ColV plasmid identified shared homology with previously reported APEC-associated plasmids, comparison of gene clusters was performed using Clinker (41).

Multiple conserved gene clusters were observed to be shared with previously reported APEC associated plasmids pAPEC-O2-211A-ColV (Assession number: CP030791.1) and pAPEC-p10_578_1 (Assession number: CP087565.1). However, extensive variation was observed between the pSAP-4026 ColV plasmid and the similarly sized pCh101 (Assession number: CP127318.1) previously identified within commensal avian *E. coli* (Figure 3). This suggests the ColV plasmid observed within this outbreak-associated lineage bears a greater similarity to those commonly found within high-risk virulent clonal groups compared to commensal avian *E. coli*.

**Figure 3.**
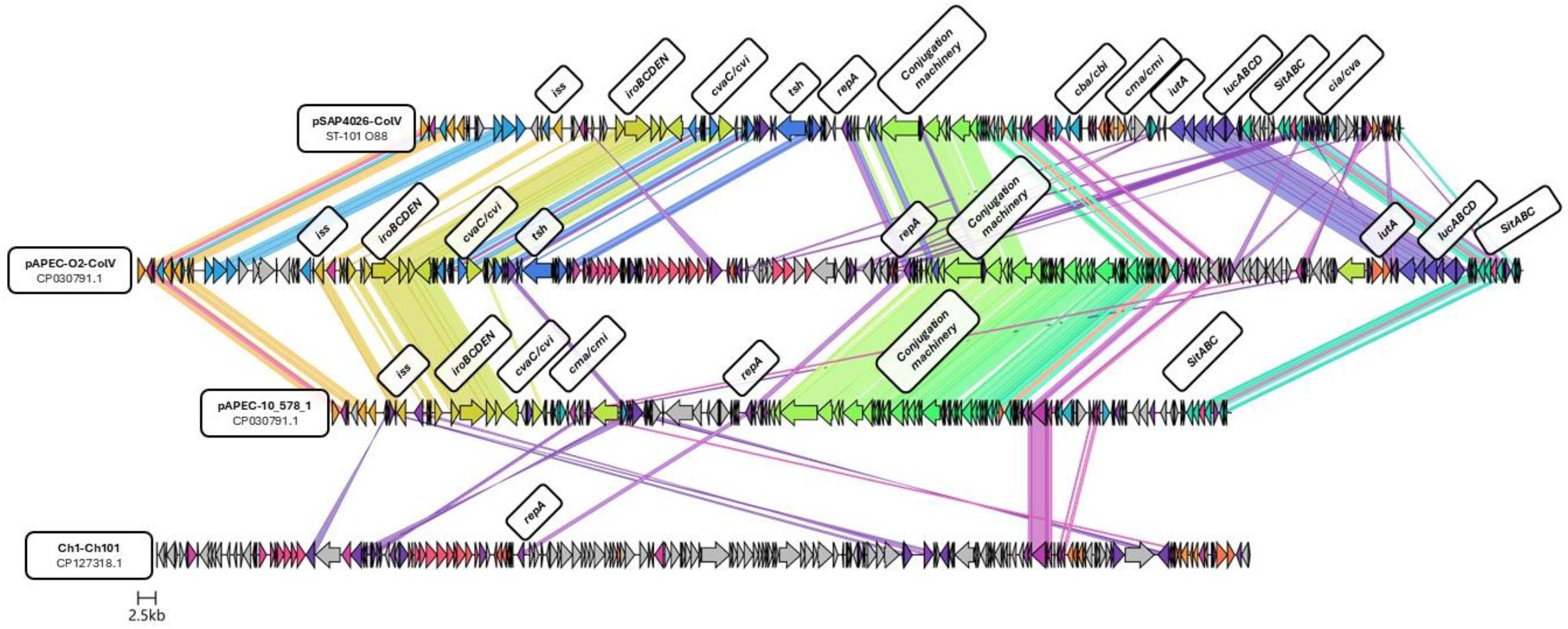
Gene cluster comparison of pSAP4026-ColV with previously reported plasmids from avian pathogenic and commensal *E. coli*. Figure generated using the clinker and cluster map software (41) with identity threshold set at 0.3 with additional labelling performed in biorender (www.biorender.com). Plasmid sequences of APEC-O2-211A-ColV (Assession number: CP030791.1), pAPEC-p10_578_1 (Assession number: CP087565.1), and pCh101 (Assession number: CP127318.1) accessed through GenBank.

Collectively, characterisation of the colibacillosis outbreak identified ST-101 O88:H8 as the dominant clonal lineage which was typically associated with the development of disease in birds and harboured multiple plasmid virulence factors.

### APEC ST-101, ST-95, and ST-140 exhibit higher levels of invasion of 8E11 chicken gut epithelial cells than ST-23 and ST-117

Within the colibacillosis outbreak, multiple APEC genotypes were observed, however it was dominated by the ST-101 clonal group. ST-101 APEC have previously been reported within poultry in multiple countries (15, 32, 43-45), in addition to being associated with recent colibacillosis outbreaks in Denmark (46). Yet, despite the growing prominence of this lineage, the phenotypic profile of ST-101 and how it may differ from other high-risk clonal groups remains poorly understood. Therefore, ST-101 APEC isolated from this outbreak in turkeys was compared to predominant APEC lineages frequently associated with avian colibacillosis (Table 1) using the 8E11 chicken gut epithelium *in vitro* model. Adhesion to cellular surfaces prevents mechanical clearance and confers a selective advantage for pathogenic bacteria (47). Here, all APEC isolates demonstrated the ability to adhere to the 8E11 cell line, with no significant difference (*p* ≤ 0.05) observed between APEC lineages (Figure 4A). Comparison to the invasive pathogen *S*. Typhimurium saw no significant difference in adherence capacity of APEC isolates, with the exception of APEC ST-95 O18 where adherence was significantly (*p* ≥ 0.05) reduced compared to *S*. Typhimurium.

**Figure 4:**
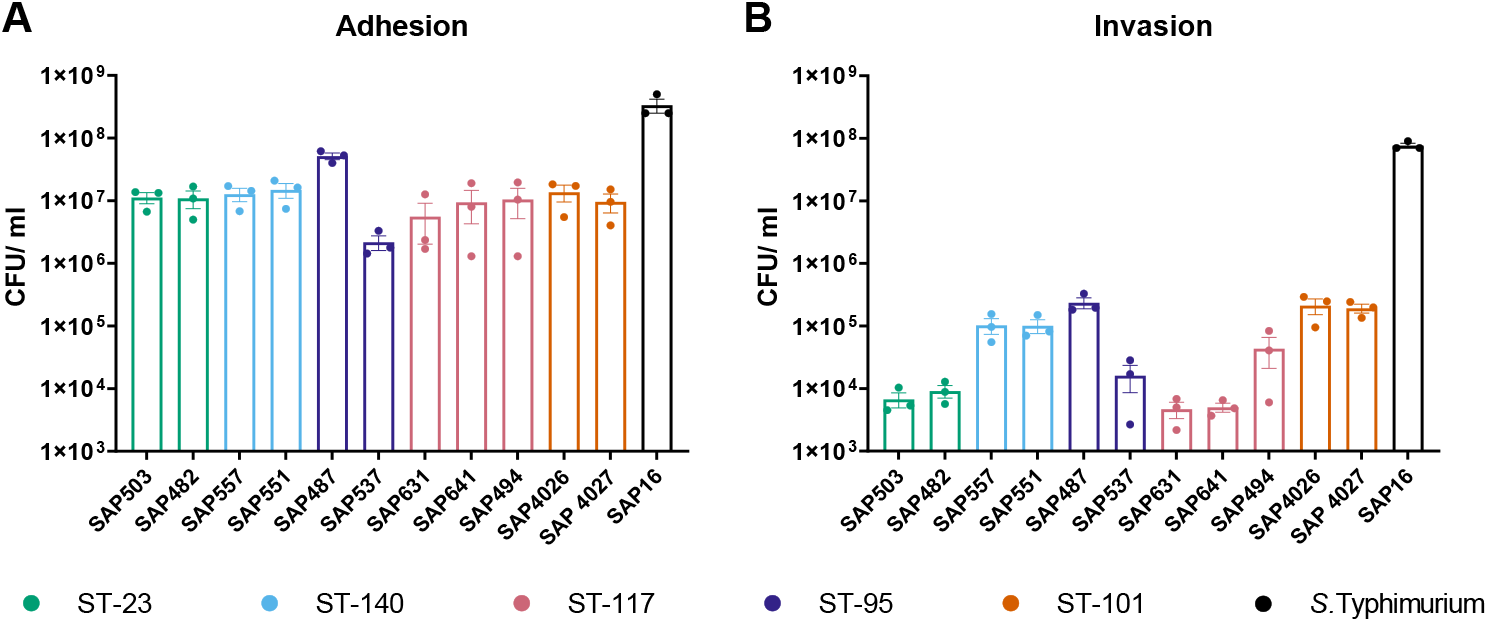
Comparison of ability of APEC isolates to (A) adhere to and (B) invade the 8E11 cell line. Quantification of adhesion of APEC isolates determined following challenge with MOI 100 bacterial inoculum followed by two hours incubation at 41°C prior to washing and lysing of eukaryotic cells. Invasion assayed by the inoculation of cells for two hours followed by the addition of cell culture media containing gentamicin for a further two hours, before washing and lysis of eukaryotic cells. Bacterial viability determined by the Miles and Misra method. Experiments performed independently three times, with triplicate experimental repeats. Data shows average CFU/ml ± SEM.

**Figure 5:**
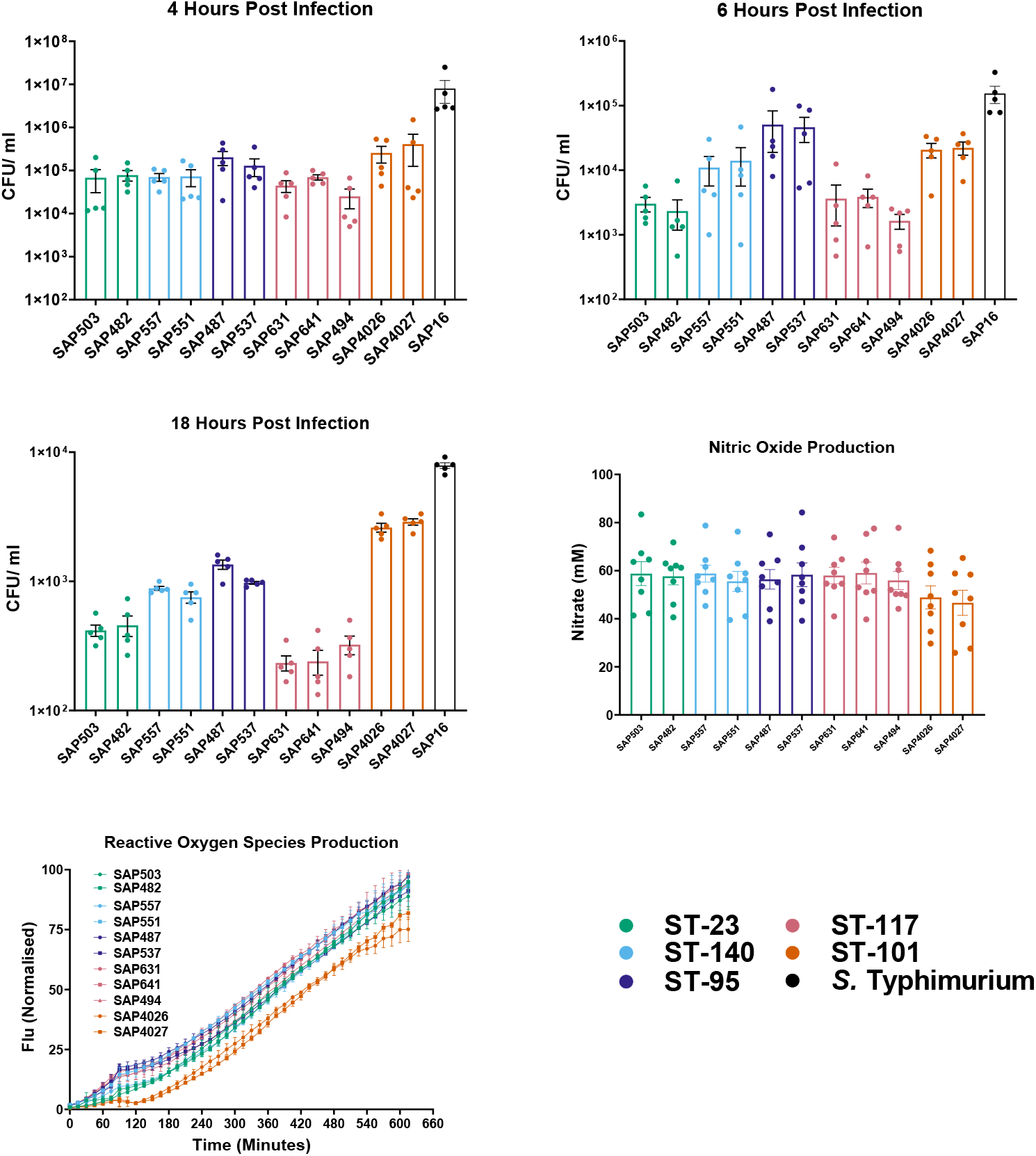
Comparison of intracellular survival of APEC isolates in HD11 chicken macrophage cells at (A) 4 hours post infection, (B) 6 hours post infection, and (C) 18 hours post infection. Quantification of intracellular survival of APEC isolates determined following challenge with MOI 10 bacterial inoculum followed by two hours incubation at 41°C prior to addition of cell culture media containing gentamicin for described incubation period. HD11 cells were then washed with PBS and eukaryotic cells lysed. Bacterial viability determined by the Miles and Misra method. Experiments performed independently five times, with triplicate technical repeats. Data shows average CFU/ml ± SEM.

In addition to adhesion, *E. coli* and other Gram-negative pathogens have been observed to invade host cells, allowing intracellular replication and reduced competition for nutrients (47, 48). Ability of APEC isolates to invade 8E11 cells showed significant (*p* = 0.0013) variation between APEC lineages (Figure 4B). Invasive ability compared to *S*. Typhimurium was also observed to be significantly lower in the ST-23 SAP503 (*p* =0.018) and SAP482 (*p* =0.065), the ST-95 isolate SAP537 (*p* =0.058), and ST-117 SAP631 and SAP 641 (*p* =0.008). In contrast, isolates belonging to ST101, ST-140, and the ST-95 SAP487 isolate demonstrated a greater degree of invasion, though still lower than *S*. Typhimurium (*p* > 0.999). Intra-sequence type variations were also observed within isolates with different O-antigen serotypes, with SAP487 O45 and SAP537 O18 demonstrating both the highest and lowest invasive ability, respectively. All APEC lineages demonstrated similar (*p* ≤ 0.05) growth rate in 8E11 cell culture media (Figure S1), indicating that differences in bacterial load recovered was not a result of increased growth. Collectively, this suggests that genotype defines the capacity of APEC to invade gut epithelial cells.

### ST-101 APEC demonstrated increased intracellular survival within HD11 chicken macrophage cells compared to ST-117 APEC

Macrophages are the first line of defence following infection, phagocytose and killing invading pathogens. However, many pathogens, including some *E. coli* (49), have also evolved to resist phagocytosis and intracellular killing by macrophages, thereby aiding systemic infection. The *in vitro* HD11 avian macrophage cell line was used to compare the ability of APEC lineages to survive intracellularly. At four hours post infection, all APEC sequence types exhibited comparable (*p* =0.1308) intracellular survival within chicken macrophages (Figure 6A). However, significant (*p* <0.001) differences between APEC sequence types were present at six hours (Figure 6B) and 18 hours post infection (Figure 6C). Comparison of intracellular survival to that of *S*. Typhimurium revealed trends between sequence types, alike those observed within 8E11 cell invasion (Figure 4). At both six hours and 18 hours post infection, bacterial survival was significantly reduced in ST-23 isolates SAP503 (*p* >0.001) and SAP482 (*p* >0.01), as well as ST117 isolates SAP631 (*p* >0.01), SAP641 (*p* >0.001), and SAP494 (*p* >0.001) compared to *S*. Typhimurium. Meanwhile, isolates belonging to ST-140, ST-95, and ST-101 demonstrated high intracellular survival, comparable (*p* ≤ 0.05) to *S*. Typhimurium. Adhesion to HD11 chicken macrophage was confirmed to be equivalent (*p* ≤ 0.05) between APEC isolates tested, suggesting that changes in intracellular survival were not a result of reduced phagocytosis (Figure S2).

**Figure 6:**
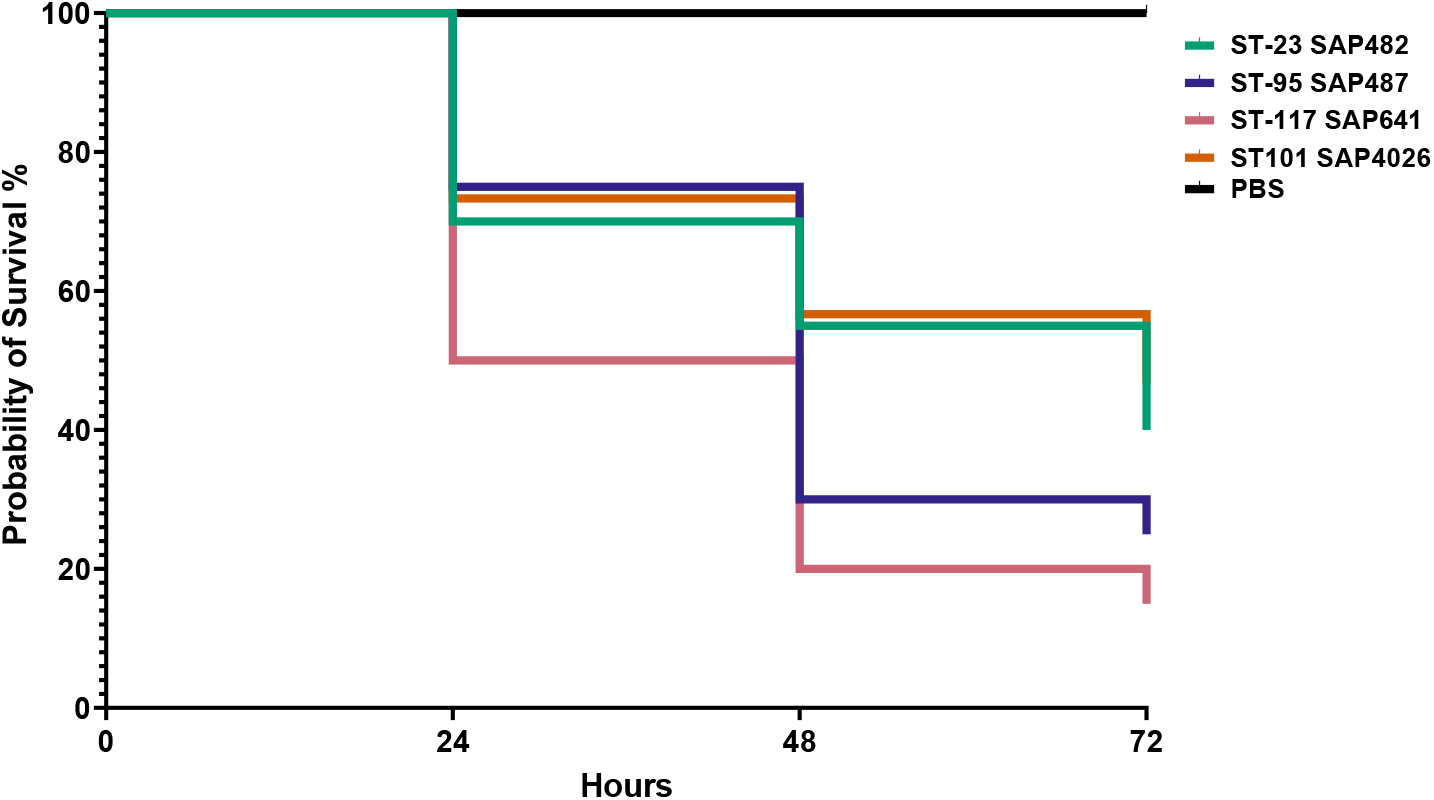
Survival of *Galleria mellonella* challenged with APEC clonal groups ST-23, ST-95, ST-117, and ST-101 24 hours, 48 hours, and 72 hours following challenge, expressed by Kaplan-Meier plots. Larvae challenged with 1×10^3^ CFU APEC or PBS mock and survival determined every 24 hours by confirming response to stimuli. Experiment performed with 10 larvae per group. Significance determined log-rank (Mantel-Cox) test.

Within macrophages, nitric oxide (NO) and reactive oxygen species (ROS) are frequently used as an indicator of the magnitude of inflammatory immune response, functioning as antimicrobial agents. Nitric oxide production was induced by all isolates following exposure and incubation for 18 hours, with no significant (*p* ≤ 0.05) differences observed (Figure S1). Similarly, ROS production was induced following infection of HD11 cells with all APEC lineages (Figure S2). Macrophages infected with ST-101 APEC exhibited delayed production of ROS compared to other APEC lineages, but there was no significant (*p* ≤ 0.05) difference in ROS production between the isolates overall. Together, this is indicative that APEC clonal lineages have differing capacity to survive within avian macrophages, despite comparable immunological responses following infection.

### Virulence in *Galleria mellonella* is varied between APEC clonal groups

The *Galleria mellonella* infection model has been used frequently as an indicator of microbial virulence (38, 39, 50, 51), and has recently been demonstrated to be a viable model for determining virulence in APEC, correlating with virulence assays performed in one-day old chicks (52). The larval innate immune system has functional similarities to that of vertebrates (53), allowing investigation of innate immune cell-pathogen interactions. *Galleria* larvae were used to compare the virulence of APEC clonal groups and establish whether enhanced intracellular survival observed in ST-101 and ST-95 correlated with a greater virulence potential.

Larvae were challenged with ST-23 SAP482, ST-95 SAP487, ST-117 SAP641, and ST-101 SAP4026, with survival assessed as an indicator of virulence. Comparison of survival curves revealed a trend (*p* = 0.0537) of differing larval mortality between all APEC isolates (Figure 6). Despite demonstrating lower invasion and intramacrophage survival relative to other clonal groups, the ST-117 SAP487 isolate caused the highest mortality at each timepoint examined, significantly increased compared to ST-23 SAP482 (*p* =0.0487) and ST-101 SAP4026 (*p* = 0.011). The ST-95 SAP487 isolate also displayed a unique virulence trend, with increased mortality at 48 hours relative to SAP482 and SAP503. Collectively, this suggests that the virulence of APEC in a *Galleria* infection model differs between APEC clonal groups and may not be dependent on intracellular survival.

## Discussion

The APEC pathotype is highly diverse, with several genotypically distinct high-risk clonal groups identified (12, 29). Although these groups have been extensively characterised (12, 15, 33, 34, 54-57), how these genotypes may differ from one another phenotypically and manifest disease remains poorly understood. Here, we aimed to phenotypically compare high-risk and outbreak-associated APEC clonal groups.

Genomic characterisation of APEC from a colibacillosis outbreak in turkey poults identified multiple genotypes associated with disease, however ST-101 was observed to be the dominant outbreak lineage, consistent with previous reports (46). ST-101 isolates encoded a high density of virulence genes on a ColV plasmid which shares large regions of homology with previously reported APEC plasmids. Investigation of adhesion and invasion capacity of ST-101 APEC and major APEC lineages in the 8E11 chicken gut epithelial cell line revealed isolates belonging to ST-95, ST-140, and ST-101 possess higher invasive ability compared to ST-23 and ST-117. A similar relationship was also observed in the capacity of APEC isolates to survive intracellularly in the HD11 chicken macrophage cell line, with ST-23 and ST-117 demonstrating reduced survival compared to other sequence types examined. Infection of *Galleria mellonella* suggested that APEC lineages have varied virulence capability, and that ST-117 was the most virulent of all lineages tested despite reduced epithelial invasion and intramacrophage survival relative to other STs.

Previous studies have aimed to characterise APEC outbreaks within poultry and documented the incidence of specific AMR and virulence genes (32, 58-60), as well as the contribution to virulence by APEC associated plasmids (24, 25). Here, we identified ST-101 as an outbreak strain; this lineage has been detected globally and heavily associated with multidrug resistance (MDR) and pan-drug resistance phenotypes (61, 62), human extraintestinal infection (63-68), as well as the environment (69). Within birds, ST-101 was first identified in a colibacillosis outbreak in Spain (32), but has since been detected in Germany, Australia, Sweden, Brazil, and the UK (15, 43-45). It is therefore imperative the characteristics of this lineage are understood so that it can be managed effectively.

We have previously described the predominant APEC high-risk clonal groups ST-23 (phylogroup C) and ST-117 (phylogroup G) APEC are commonly reported in colibacillosis outbreaks (30-32), and comprise the most reported serotype in APEC infection, O78 (12). Additionally, ST-95 and ST-140 APEC comprise two of the three distinct subpopulations within APEC-associated phylogroup B2 (12, 33). Within turkeys, the majority of clinical isolates belong to phylogroup B2, while caecal isolates are predominantly phylogroup B1 (29). With ST-101 identified as belonging to phylogroup B1, this may suggest that this lineage may have a distinct evolutionary trajectory leading to a unique mechanism of action and eventual capacity to cause disease within birds. Therefore, cell culture models were utilised to investigate the mechanism of action of the ST-101 lineages, in comparison to high-risk clonal APEC clonal groups. Here, APEC sequence types were observed to differ in their invasive capacity. Invasion of host cells can provide several advantages to bacterial pathogens, providing access to nutrients and providing protection from immunological responses (47). All lineages investigated demonstrated the ability to adhere to, and invade the gut epithelial cell line, a phenotype commonly associated with most *E. coli* pathotypes through the action of surface binding proteins (70). Isolates belonging to ST-95, ST-140, and ST-101 had comparatively higher levels of invasion within the 8E11 cell line relative to ST-23 and ST-117. The invasive ability of APEC has been previously investigated in 8E11 cells (71), as well as tracheal (72), lung (73, 74), and liver epithelial cells (74), wherein invasion has been reported, mirroring what has been observed within this study. However, due to vast difference in cell type and the use of only a single APEC isolate, it is difficult to corroborate the observed differences between sequence type. However, comparison of an ST-101 O154 ExPEC isolated from a human patient to other ExPEC from phylogroup B2 highlighted an enhanced ability to invade epithelial in ST-101 ExPEC (62). This contrasts with observations made in this study, where ST-101 APEC had similar invasive ability to isolates belonging to phylogroup B2 (ST-95 and ST-140). This discordance may be a result of differing virulence factors present between APEC clonal groups, with pronounced virulence gene profiles observed within previously characterised colibacillosis outbreaks (15, 16), as well as APEC characterised within this study (Figure 1).

In addition to the invasion of gut epithelial cells, the ability of the representative APEC lineages to survive within the HD11 chicken macrophage cell line was examined. *E. coli* have been frequently demonstrated to resist macrophage killing and even proliferate following phagocytosis (49, 75-80). The ability of APEC to persist within HD11 cells has also been previously reported (81), similarly all APEC demonstrated the ability to survive at comparably low cell density beyond 18 hours post infection, but trends between intramacrophage survival and sequence type were observed. Alike the invasion of 8E11 cells, ST-95, ST101, and ST-140 demonstrated increased intracellular survival relative to ST-23 and ST-117. Enhanced intramacrophage survival of ST-140 and ST-95 isolates may be a result of the presence of the K1 capsule, typically associated with phylogroup B2 *E. coli* (82). K1 capsules have been observed to confer resistance to phagocyte engulfment and killing in *E. coli* (83), Group B *Streptococcus* (84), and *Staphylococcus aureus* (85). Previous studied have also investigated the interactions of APEC strains with phagocytes, with Mellata *et al*. observing reduced association with, and phagocytic bactericidal activity against, serotype O78 APEC (assumed to be ST-23 or ST-117) relative to O1 or O2 APEC in primary macrophages and heterophils (86). In contrast, no reduction in association was observed within this study with O78 APEC, in addition to increased bacterial clearance. This may be a result of differing genomic background between isolates, opposed to O-antigen identity, with Mellata *et al*. also observing increased type 1 fimbriae expression enhancing resistance of phagocyte killing. Furthermore, additional studies have observed a continuous reduction in O78 APEC intramacrophage survival (81), while other work showed an O2 isolate had improved survival and demonstrated the ability to replicate within the same cell line (87).

The *Galleria mellonella* infection model has been used frequently as an indicator of microbial virulence (38, 39, 50, 51), and has recently been demonstrated to be a viable model for determining virulence in APEC, correlating with virulence assays performed in one-day old chicks (52). The larval innate immune system also has functional similarities to that of vertebrates (53), allowing investigation of innate immune cell-pathogen interactions. Moreover, previous investigations have shown that virulence of ExPEC isolates within the *Galleria mellonella* infection model differs between sequence types (88, 89). Similarly, this study presents differences in virulence potential between APEC sequence types, with SAP641 ST-117 showing greatest virulence potential. ST-117 APEC are belonging to phylogroup G have recently emerged as the most frequently implicated cause of colibacillosis and are associated with high pathogenicity (6, 29, 59, 90). Therefore, it is unsurprising that this lineage displays high levels of virulence with *Galleria* larvae. This high *in vivo* virulence potential is not consistent with observed phenotypic interactions within the *in vitro* 8E11 and HD11 cell culture models, wherein invasion and intramacrophage survival of ST-117 was low relative to ST-101 isolates. These findings make clear that the capacity to invade and survive intracellularly are not the only process that dictate virulence potential within APEC. Moreover, it is suggestive of distinct routes of pathogenesis and host cell interactions between APEC clonal groups. Further investigation characterising the phenotypic nature of APEC genotypes is necessary to elucidate the determinants of these distinct behaviours outside of epithelial and macrophage interactions, as modelled with *Galleria mellonella* and how this may be combated with protective therapies.

In summary, we present evidence to suggest that pronounced differences in phenotypic interactions with host cells and virulence are present between high-risk APEC clonal groups. This provides further evidence that the genotypes most commonly implicated in colibacillosis exhibit considerable heterogeneity with regards to their pathogenic potential. Future investigations into the genetic determinants that govern phenotypic variation between these genotypes is essential to inform potential future disease management strategies.

## Author contributions

Conceptualisation, J.R.G.A, S.B, R.M.L.R, and J.W.M,; Formal analysis, J.R.G.A, H.L, C.B, K.Q, S.P, K.W, E.K and J.W.M,; Funding acquisition R.M.L.R, S.B, and J.W.M,; Methodology J.R.G.A, S.B, R.M.L.R, and J.W.M,; Writing-original draft J.R.G.A, and J.W.M,; Writing-review and editing J.R.G.A, K.W, S.B, R.M.L.R, and J.W.M.

## Data Availability

The authors confirm all raw data and supplementary files have been provided within the article or are available in the public repository Zenodo DOI: 10.5281/zenodo.14965605. Genome sequence data are available within the Genomes repository at NCBI, in Bioproject Accession PRJNA1222627, and available from Zenodo DOI: 10.5281/zenodo.14892864. The information for the APEC genomes has been included in Supplementary File 1.

## Funding information

BB/T020954/1; <u>Biotechnology and Biological Sciences Research Council/United Kingdom</u>

BB/T020954/2; <u>Biotechnology and Biological Sciences Research Council/United Kingdom</u>

BB/S01506X/1; <u>Biotechnology and Biological Sciences Research Council/United Kingdom</u>

BBS/E/PI/230001A; <u>Biotechnology and Biological Sciences Research Council/United Kingdom</u>

The British Egg Marketing Research and Education Trust, United Kingdom

## Ethical approval

Favourable ethical opinion for bacterial isolate collection was confirmed by the University of Surrey Non Animal (Scientific Procedures) Act (NASPA) sub-committee of the Animal Welfare and Ethical Review Board (NASPA-2424-09(L)). This study has adhered to ARRIVE guidelines.

## Consent for publication

All authors consent for the publication of this study.

## Acknowledgements

The authors acknowledge the BBSRC and The British Egg Marketing Research and Education Trust for funding this study. Additionally the authors gratefully acknowledge the participating farms and Avara Foods Ltd for providing bacterial isolates. We would also like to acknowledge MicrobesNG who provided the genome sequencing service (http://www.microbesng.uk).

