## Supplementary File for "High-risk clonal groups of Avian Pathogenic *Escherichia coli* (APEC) demonstrate heterogeneous phenotypic characteristics *in vitro* and *in vivo*"

### Supplementary material

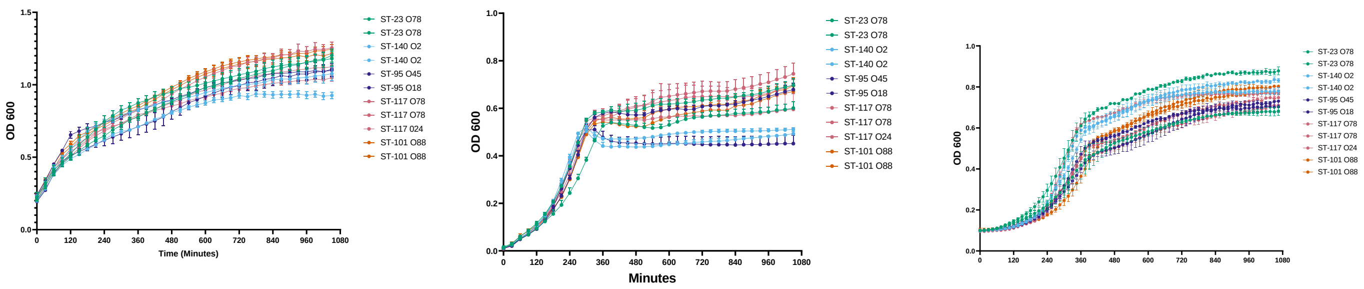


**Figure S1** Microbial growth curves of APEC isolates belonging to distinct sequence types in (A) LB broth, (B) DMEM media supplemented with 10% FBS, and (C) RPMI media supplemented with 5% FBS and 5% CS. Inoculated media was incubated at 37°C for 18 hours, aerobically with optical density at 600 nm recorded every 30 minutes using a TECAN plate reader. Data represents three independent experiments performed in triplicate and presented as mean $OD_{600}$ ±SEM.

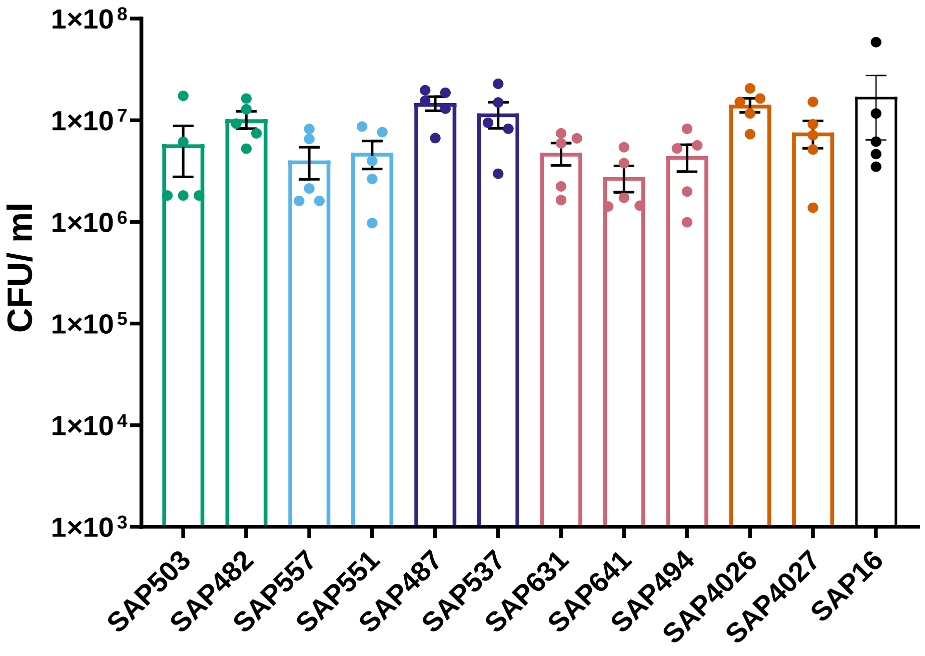


**Figure S2** Comparison of adhesion of APEC isolates in HD11 chicken macrophage cells. Quantification of adhesion APEC isolates determined following challenge with MOI 10 bacterial inoculum followed by two hours incubation at 41°C prior to washing with PBS and lysis. Bacterial viability determined by the Miles and Misra method.

Experiments performed independently five times, with triplicate experimental repeats. Data shows average CFU/ml ± SEM.

Table S1

| **APEC isolate name** | **BioSample Accession** | **Collection Year** | **Source** |
|---|---|---|---|
| pSAP4027 | SAMN46824902 | 2020 | Turkey |
| SAP4026 | SAMN46824900 | 2020 | Turkey |
| SAP4027 | SAMN46824901 | 2020 | Turkey |
| SAP4442 | SAMN46824810 | 2020 | Turkey |
| SAP4443 | SAMN46824811 | 2020 | Turkey |
| SAP4444 | SAMN46824812 | 2020 | Turkey |
| SAP4445 | SAMN46824813 | 2020 | Turkey |
| SAP4446 | SAMN46824814 | 2020 | Turkey |
| SAP4447 | SAMN46824815 | 2020 | Turkey |
| SAP4448 | SAMN46824816 | 2020 | Turkey |
| SAP4449 | SAMN46824817 | 2020 | Turkey |
| SAP4450 | SAMN46824818 | 2020 | Turkey |
| SAP4451 | SAMN46824819 | 2020 | Turkey |
| SAP4452 | SAMN46824820 | 2020 | Turkey |
| SAP4453 | SAMN46824821 | 2020 | Turkey |
| SAP4454 | SAMN46824822 | 2020 | Turkey |
| SAP4455 | SAMN46824823 | 2020 | Turkey |
| SAP4456 | SAMN46824824 | 2020 | Turkey |
| SAP4457 | SAMN46824825 | 2020 | Turkey |
| SAP4458 | SAMN46824826 | 2020 | Turkey |
| SAP4459 | SAMN46824827 | 2020 | Turkey |
| SAP4460 | SAMN46824828 | 2020 | Turkey |
| SAP4461 | SAMN46824829 | 2020 | Turkey |
| SAP4462 | SAMN46824830 | 2020 | Turkey |
| SAP4463 | SAMN46824831 | 2020 | Turkey |
| SAP4464 | SAMN46824832 | 2020 | Turkey |
| SAP4465 | SAMN46824833 | 2020 | Turkey |
| SAP4466 | SAMN46824834 | 2020 | Turkey |
| SAP4467 | SAMN46824835 | 2020 | Turkey |
| SAP4468 | SAMN46824836 | 2020 | Turkey |
| SAP4469 | SAMN46824837 | 2020 | Turkey |
| SAP4470 | SAMN46824838 | 2020 | Turkey |
| SAP4471 | SAMN46824839 | 2020 | Turkey |
| SAP4472 | SAMN46824840 | 2020 | Turkey |
| SAP4473 | SAMN46824841 | 2020 | Turkey |
| SAP4474 | SAMN46824842 | 2020 | Turkey |
| SAP4475 | SAMN46824843 | 2020 | Turkey |
| SAP4476 | SAMN46824844 | 2020 | Turkey |

| | | | |
|---|---|---|---|
| SAP4477 | SAMN46824845 | 2020 | Turkey |
| SAP4478 | SAMN46824846 | 2020 | Turkey |
| SAP4479 | SAMN46824847 | 2020 | Turkey |
| SAP4480 | SAMN46824848 | 2020 | Turkey |
| SAP4481 | SAMN46824849 | 2020 | Turkey |
| SAP4482 | SAMN46824850 | 2020 | Turkey |
| SAP4483 | SAMN46824851 | 2020 | Turkey |
| SAP4484 | SAMN46824852 | 2020 | Turkey |
| SAP4485 | SAMN46824853 | 2020 | Turkey |
| SAP4486 | SAMN46824854 | 2020 | Turkey |
| SAP4487 | SAMN46824855 | 2020 | Turkey |
| SAP4488 | SAMN46824856 | 2020 | Turkey |
| SAP4489 | SAMN46824857 | 2020 | Turkey |
| SAP4490 | SAMN46824858 | 2020 | Turkey |
| SAP4491 | SAMN46824859 | 2020 | Turkey |
| SAP4492 | SAMN46824860 | 2020 | Turkey |
| SAP4493 | SAMN46824861 | 2020 | Turkey |
| SAP4494 | SAMN46824862 | 2020 | Turkey |
| SAP4495 | SAMN46824863 | 2020 | Turkey |
| SAP4496 | SAMN46824864 | 2020 | Turkey |
| SAP4497 | SAMN46824865 | 2020 | Turkey |
| SAP4498 | SAMN46824866 | 2020 | Turkey |
| SAP4499 | SAMN46824867 | 2020 | Turkey |
| SAP4500 | SAMN46824868 | 2020 | Turkey |
| SAP4501 | SAMN46824869 | 2020 | Turkey |
| SAP4502 | SAMN46824870 | 2020 | Turkey |
| SAP4503 | SAMN46824871 | 2020 | Turkey |
| SAP4504 | SAMN46824872 | 2020 | Turkey |
| SAP4505 | SAMN46824873 | 2020 | Turkey |
| SAP4506 | SAMN46824874 | 2020 | Turkey |
| SAP4507 | SAMN46824875 | 2020 | Turkey |
| SAP4508 | SAMN46824876 | 2020 | Turkey |
| SAP4509 | SAMN46824877 | 2020 | Turkey |
| SAP4510 | SAMN46824878 | 2020 | Turkey |
| SAP4511 | SAMN46824879 | 2020 | Turkey |
| SAP4512 | SAMN46824880 | 2020 | Turkey |
| SAP4513 | SAMN46824881 | 2020 | Turkey |
| SAP4514 | SAMN46824882 | 2020 | Turkey |
| SAP4515 | SAMN46824883 | 2020 | Turkey |
| SAP4516 | SAMN46824884 | 2020 | Turkey |
| SAP4517 | SAMN46824885 | 2020 | Turkey |
| SAP4518 | SAMN46824886 | 2020 | Turkey |
| SAP4519 | SAMN46824887 | 2020 | Turkey |
| SAP4520 | SAMN46824888 | 2020 | Turkey |
| SAP4521 | SAMN46824889 | 2020 | Turkey |
| SAP4522 | SAMN46824890 | 2020 | Turkey |

| | | | |
|---|---|---|---|
| SAP4523 | SAMN46824891 | 2020 | Turkey |
| SAP4524 | SAMN46824892 | 2020 | Turkey |
| SAP4525 | SAMN46824893 | 2020 | Turkey |
| SAP4526 | SAMN46824894 | 2020 | Turkey |
| SAP4527 | SAMN46824895 | 2020 | Turkey |
| SAP4528 | SAMN46824896 | 2020 | Turkey |
| SAP4529 | SAMN46824897 | 2020 | Turkey |
| SAP4530 | SAMN46824898 | 2020 | Turkey |
| SAP4531 | SAMN46824899 | 2020 | Turkey |
